# Activity and Dynamics of p110α are not Differentially Modulated by Regulatory Subunit Isoforms

**DOI:** 10.1101/2025.10.16.682970

**Authors:** Isobel Barlow-Busch, Emma E Walsh, Hunter G Nyvall, John E Burke

## Abstract

Class IA phosophoinositide kinases (PI3Ks) are master regulators of growth, metabolism, and immunity. The class IA PI3Ks are a heterodimer composed of a p110 catalytic subunit and one of five possible regulatory subunits (p85α, p85β, p55γ, p55α, p50α). The regulatory subunit plays critical roles in stability, inhibition, and activation of the p110 catalytic subunit. The p110α catalytic subunit frequently contains activating mutations in human cancer, with many of these mutations altering the interaction between catalytic and regulatory subunits. It has been found that different regulatory subunits play unique roles in human disease, but it is unknown how these different subunits regulate p110α. Here, using a synergy of biochemical assays and hydrogen deuterium exchange mass spectrometry (HDX-MS) we examined how the five different regulatory subunits inhibit, activate, and interact with the p110α catalytic subunit. We find that there are no significant differences in lipid kinase activity or in membrane recruitment between the different heterodimer complexes. HDX-MS in the presence and absence of an activating phosphopeptide also showed only minor conformational differences between different regulatory subunit complexes. Overall, our work reveals that the different regulatory subunits interact with and inhibit p110α in a similar fashion at a molecular level.

## Introduction

The class I phosphoinositide 3-kinases (PI3Ks) are crucial lipid signaling enzymes responsible for generating the lipid second messenger phosphatidylinositol (3,4,5)-trisphosphate (PIP_3_) downstream of various cell surface receptors, including receptor tyrosine kinases (RTKs), G-protein coupled receptors (GPCRs), and Ras (Burke, 2018; Burke and Williams, 2015). PIP_3_ subsequently activates pro-growth signaling events, most notably the protein kinase Akt, which drives processes such as cell growth, proliferation, survival, and nutrient uptake (Madsen and Vanhaesebroeck, 2020; Manning and Toker, 2017; Shaw and Burke, 2025). Class I PI3Ks are divided into two subgroups, class IA composed of three catalytic subunits (p110α, p110β, p110γ) and class IB composed of a single p110γ catalytic subunit (Shaw et al., 2025). The main difference between class IA and class IB is the association with different regulatory subunits, with class IA having five regulatory subunits (p85α, p55α, p50α, p85β, and p55γ) and class IB having two (p101 and p84) (Rathinaswamy and Burke, 2019). PI3K complexes are frequently referred to by the identity of the catalytic subunit, with PI3Kα referring to a complex containing p110α with any possible regulatory subunit.

PI3Kα exists as a hetero-dimer composed of a p110α catalytic subunit and a regulatory subunit. The p110α catalytic subunit is a large multi-domain protein consisting of an adaptor-binding domain (ABD), a Ras-binding domain (RBD), a C2 domain, a helical domain, and a bilobal kinase domain (Figure 1A) (Huang et al., 2007; Miled et al., 2007). The regulatory subunits (p85α, p85β, p55α, p50α, and p55γ) are encoded by three distinct genes (*PIK3R1, PIK3R2*, and *PIK3R3*) and share a core architecture comprising an N-terminal Src homology 2 (nSH2) domain, an inter-SH2 (iSH2) coiled-coil domain, and a C-terminal Src homology 2 (cSH2) domain (Miled et al., 2007; Piccione et al., 1993). The two longest isoforms, p85α (*PIK3R1*) and p85β (*PIK3R2*), contain the complete set of domains: an N-terminal Src homology 3 (SH3) domain and a Bcr homology (BH) domain flanked by proline-rich motifs. Although the C-terminal regions (nSH2–iSH2–cSH2) of p85α and p85β are highly homologous (∼75% identity), their N-terminal regions (SH3– BH) are more divergent, showing only ∼37% identity(Vallejo-Díaz et al., 2019).

**Fig 1.**
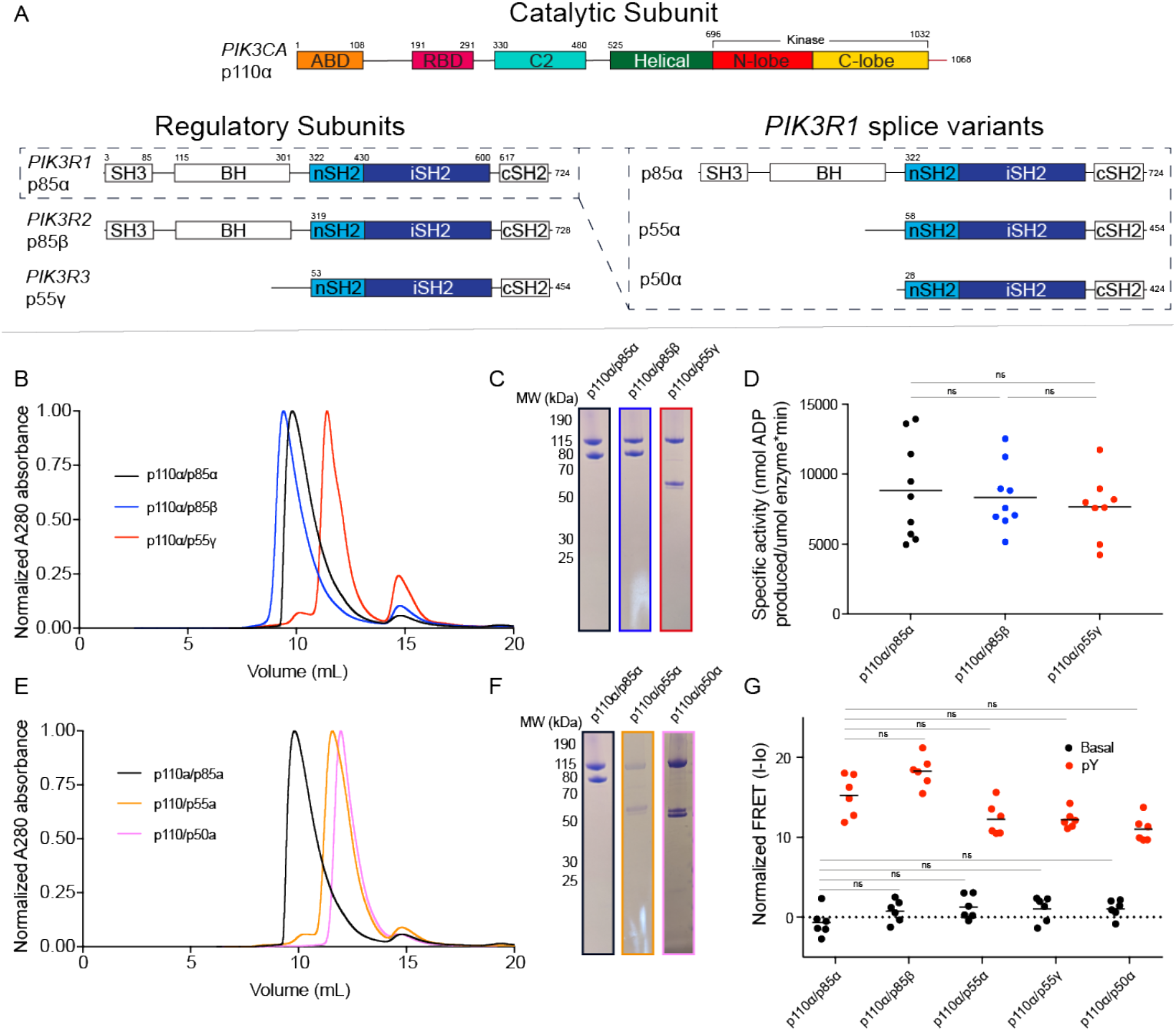
Domain architecture, purification, and biochemical characterization of PI3Ka complexes. **A**. Domain architecture of the catalytic subunit p110α (encoded by *PIK3CA*) and the regulatory subunits p85α, p85β, and p55γ (encoded by *PIK3R1, PIK3R2*, and *PIK3R3*). *PIK3R1* has three splice variants, including p55α and p50α, which lack the N-terminal SH3 and BH domains. **B**. Gel filtration traces of p110α/p85α, p110α/p85β, and p110α/p55γ, with size standards indicated. **C**. SDS–PAGE gel of purified p110α/p85α, p110α/p85β, and p110α/p55γ, with size standards indicated. **D**. Specific activity of PI3Kα complexes p110α/p85α, p110α/p85β, and p110α/p55γ (technical replicates; mean values shown; n = 9 for p110α/p85α and p110α/p85β, n = 8 for p110α/p55γ). Assays were performed with 5 nM PI3K and 100 μM ATP with 0.45 mg/mL 5% PIP2/95% PS vesicles. Two-tailed t-test results: ns, p > 0.05. **E**. Gel filtration traces of p110α/p85α, p110α/p55α, and p110α/p50α, with size standards indicated. **F**. SDS–PAGE gel of purified p110α/p85α, p110α/p55α, and p110α/p50α, with size standards indicated. **G**. Protein–lipid FRET measurements of membrane recruitment for p110α/p85α, p110α/p85β, p110α/p55γ, p110α/p55α, and p110α/p50α under basal and pY-activated conditions on liposomes containing 5% brain PIP2, 60% egg yolk PE, 25% brain PS, and 10% dansyl-PS (n = 6). Experiments were performed with 1 μM pY peptide, 250 nM PI3K, and 16.65 μg lipid vesicles. Values were normalized to WT apo. Two-tailed t-test results: ns, p > 0.05.

The remaining three isoforms are progressively truncated at the N-terminus. p55γ (*PIK3R3*) lacks the SH3 domain, the first proline-rich motif, and the BH domain, instead featuring a unique 30-residue N-terminal sequence followed by a conserved proline-rich motif (Pons et al., 1995). The splice variants p55α and p50α (*PIK3R1*) also lack the SH3 and BH domains but carry a unique 34-amino acid and 6-amino acid sequence, respectively, at their N-termini (Inukai et al., 1997). These differences in N-terminal domain composition between regulatory subunits are functionally significant, as these regions primarily mediate p110-independent adaptor functions, including interactions with cytoskeletal components (Fox et al., 2020).

Despite differences in sequence, all five regulatory subunits play three essential roles in modulating the p110α subunit: stabilizing p110α, inhibiting its basal lipid kinase activity, and enabling activation upon binding to pYXXM motifs found in phosphorylated receptors (Yu et al., 1998). Inhibition is mediated by several inter-and intra-subunit interfaces, including the ABD-kinase interface within p110α, the C2-iSH2 interface between subunits, and the interaction of the nSH2 domain with the C2, helical, and kinase domains (Mandelker et al., 2009; Miled et al., 2007). Full activation requires the binding of GTP-loaded Ras and the engagement of both the nSH2 and cSH2 domains by bis-phosphorylated pYXXM motifs derived from RTKs or adaptors (Buckles et al., 2017; Dornan et al., 2020; Siempelkamp et al., 2017). This engagement relieves inhibitory contacts, leading to conformational changes such as the disengagement and mobility of the ABD and p85 subunits relative to the p110α catalytic core (Burke et al., 2012; Jenkins et al., 2023).

The various class IA regulatory subunits exhibit crucial differences in their functional roles in disease, especially cancer, despite the high homology shared among their C-terminal domains. For instance, the p85α subunit (encoded by *PIK3R1*) is the most abundant isoform in normal tissues and traditionally functions as a tumor suppressor, relying on its capacity to efficiently maintain p110α in an inactive conformation (Ueki et al., 2002). Conversely, p85β (encoded by *PIK3R2*) is frequently elevated in advanced cancers and acts as a tumor driver or oncogene (Vallejo-Díaz et al., 2019) and studies suggest p85β exerts a weaker constraint on basal p110α activity compared to p85α (Cortés et al., 2012). Furthermore, p55γ (encoded by *PIK3R3*) also displays pro-tumorigenic activity (Nicolau-Neto et al., 2018; Peng et al., 2018). The contrasting roles of these highly homologous regulatory subunits, particularly in mediating PI3K activity and tumor progression, remain a significant area of investigation.

In this study, we used a combined biochemical and biophysical approach to investigate the molecular mechanisms of how regulatory subunits encoded by *PIK3R1, PIK3R2*, and *PIK3R3* influence the p110α catalytic subunit. Activity assays showed no differences in specific activity among complexes containing p85α, p85β, or p55γ. Protein-lipid FRET assays likewise revealed no differences in membrane recruitment across all five subunits, under either basal or pYXXM-activated conditions. HDX-MS analysis detected no major global conformational differences between apo-state complexes formed with subunits from the three genes, nor among *PIK3R1* splice variants (p85α, p55α, p50α) in apo or pYXXM-activated states. Together, these results suggest that the contrasting roles of regulatory subunits, such as the tumor-suppressive function of p85α versus the oncogenic role of p85β, do not arise from intrinsic biochemical differences within the p110α/regulatory subunit core complex.

## Results

### Characterization of PI3Kα Complexes with Regulatory Subunit Isoforms and Splice Variants

To enable biochemical and biophysical characterization, all five PI3Kα complexes were successfully expressed in Sf9 cells and purified. Gel filtration confirmed complex formation, with each complex eluting near its expected molecular weight (Fig. 1B). The p110α/p85β complex eluted similar to p110α/p85α whereas the p110α/p55γ, p110α /p55α, and p110α/p50α complexes eluted later, consistent with their smaller size. Purity of the constructs was verified by SDS–PAGE (Fig. 1C,E,F).

To assess intrinsic differences in catalytic activity, basal lipid kinase activity was measured using 5% PIP_2_/95% PS membranes with 5 nM PI3Kα. No statistically significant differences in specific activity were observed among complexes formed with p85α, p85β, or p55γ (Fig. 1D). These results suggest that variations in regulatory subunit sequence, or the presence/absence of N-terminal domains such as SH3 and BH, do not measurably alter the basal activity of the p110α catalytic core.

To probe differences in membrane association, we performed protein–lipid FRET assays. Membrane binding affinity of all five complexes was measured under basal and pYXXM-activated (bound to bisphosphorylated residues 735-767 of the PDGFR receptor) conditions using membranes composed of 5% PIP_2_, 10% dansyl-PS, 20% PS, and 65% PE. Upon addition of pY peptide, all complexes exhibited >10-fold activation relative to their basal state. No significant differences in membrane recruitment were observed between complexes in either basal or pY-activated conditions (Fig. 1G). A general trend of increased recruitment for full-length p85α and p85β relative to shorter isoforms (p55α, p55γ, p50α) was noted, though this was not statistically significant.

### HDX-MS Reveals Minimal Conformational Differences in the p110α Catalytic Subunit Between Regulatory Subunit Isoforms

To explore conformational effects not captured by biochemical assays, we performed HDX-MS on PI3Kα complexes. HDX-MS is a powerful technique that has been extensively used to probe the dynamics of PI3K complexes, both in solution and on membranes (Burke et al., 2012, 2011; Burke and Williams, 2013; Dornan et al., 2020, 2017; Gong et al., 2023; Harris et al., 2023; Jenkins et al., 2023; Rathinaswamy et al., 2023, 2021b, 2021c, 2021a; Walser et al., 2013). The method measures the exchange rate of amide hydrogens and acts as a surrogate for protein conformational dynamics (Masson et al., 2019, 2017).

For isoform comparisons, HDX-MS was first performed on p110α in complex with p85α, p85β, or p55γ. Exchange was monitored at 3, 30, and 3000 s (18 °C), with peptides covering 82.6% of the exchangeable amides of p110α (stats table and raw data in Source Data). Only two peptides showed significant differences (criteria: ≥0.45 Da, ≥5%, two-tailed p < 0.01) (Fig. 2A). Peptide 24–34 in the Adaptor Binding Domain (ABD), which contacts the regulatory iSH2 domain, showed decreased exchange at 3 s and increased exchange at 3000 s in the p85β complex relative to p85α (Fig. 2D). Peptide 619–632 in the helical domain showed a small increase in exchange in the p55γ complex compared to p85β (Fig. 2A,D). These differences were minor, localized, and are not in contact with the inhibitory nSH2 domain; this indicates that different regulatory subunit isoforms do not induce substantial global conformational changes in p110α at regulatory subunit interfaces.

**Fig 2.**
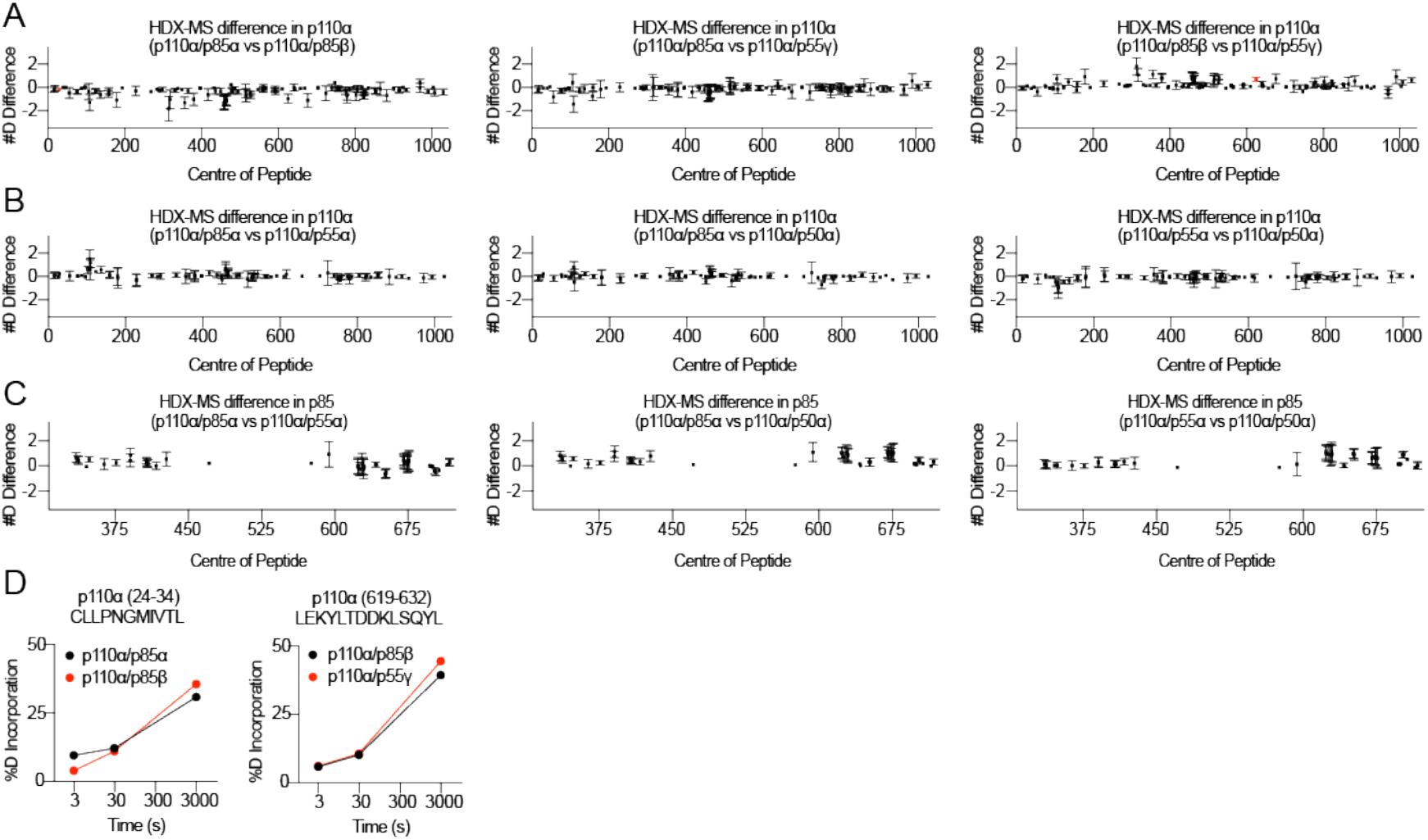
HDX-MS comparing the apo state of each p110α/regulatory subunit complex. **A**. Sum of the number of deuteron difference for all peptides in p110α analyzed across the deuterium exchange time course in p110α complexes: p110α/p85α vs p110α/p85β, p110α/p85α vs p110α/p55γ, and p110α/p85β vs p110α/p55γ. Peptides with significant changes (>0.45 Da and >5% difference at any time point, two-tailed t-test p < 0.01) are colored red. Each point represents one peptide. Error bars represent the summed S.D. across all time points (n = 3 for p110α/p85α and p110α/p85β; n = 2 for p110α/p55γ). **B**. Sum of the number of deuteron difference for all peptides in p110α analyzed across the deuterium exchange time course in p110α complexes: p110α/p85α vs p110α/p55α, p110α/p85α vs p110α/p50α, and p110α/p55α vs p110α/p50α. Significance thresholds as in (A). Each point represents one peptide. Error bars represent summed S.D. (n = 3 for each). **C**. Sum of the number of deuteron difference for all peptides spanning residues 307–724 of p85α (shared between splice variants) for the same comparisons as in (B). Significance thresholds as in (A). Each point represents one peptide. Error bars represent summed S.D. (n = 3 for each). **D**. Time courses of deuterium incorporation for two p110α peptides showing significant exchange differences between complexes with p85α, p85β, and p55γ. Means ± S.D. are shown (n = 3). Source data is provided in the Source Data file.

For splice variant comparisons, HDX-MS was performed on p110α in complex with p85α, p55α, or p50α. Identical sequences across residues 307–724 of p85α with p55α and p50α enabled direct comparisons within this region of the regulatory subunit possible. Exchange was monitored at 3, 30, and 300 s (18 °C), achieving 70.4% coverage of p110α and 67% coverage of the shared nSH2, iSH2, and cSH2 domains of the regulatory subunit. No peptides in either p110α or the regulatory subunits exhibited statistically significant differences in deuterium uptake (Fig. 2B–C) between different isoforms. These results suggest that the variable N-terminal domains (SH3, BH) of *PIK3R1* splice variants do not interact detectably with the catalytic subunit or alter the dynamics of the regulatory subunit.

### HDX-MS Reveals Minimal Conformational Differences Between Splice Variants Upon pY Activation

We next investigated whether regulatory subunit isoforms influence p110α conformational dynamics upon pY-mediated activation. HDX-MS was performed on splice variant complexes with p85α, p55α, or p50α under apo and pY-activated conditions in the absence of lipids. A schematic of pY activation is shown in Fig. 3A, highlighting disruption of the p85 nSH2 inhibitory contact sites from p110α and destabilization of the ABD–iSH2 from the catalytic core.

**Fig 3.**
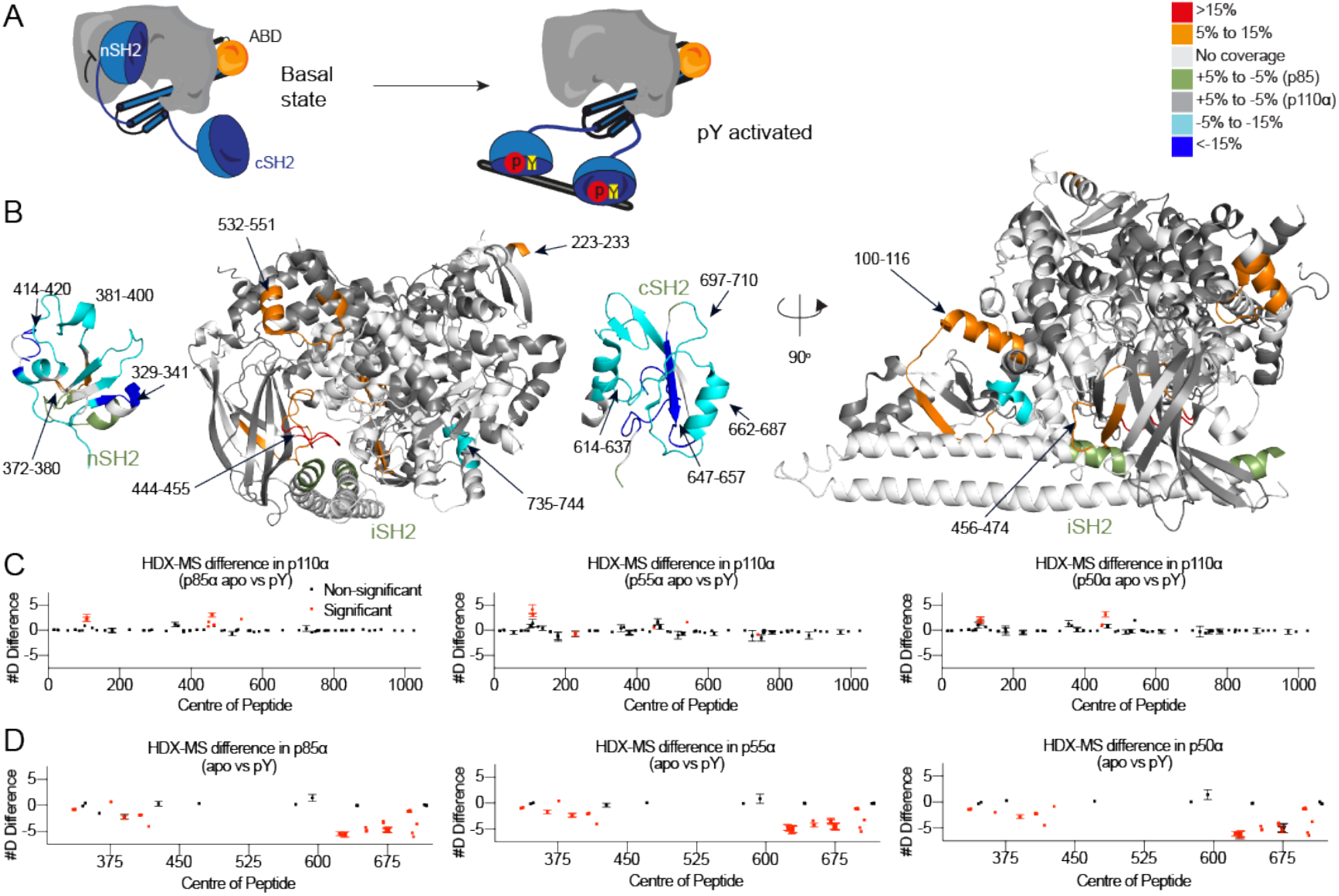
HDX-MS comparison of PI3Ka in complex with the splice variant regulatory subunits under apo and pY-activating conditions. **A**. Schematic illustrating conformational changes in PI3Kα upon pY-peptide binding, including release of the nSH2 domain. **B**. Significant differences in deuterium exchange mapped onto the PI3Kα structure. Models were built from p110α from PI3Kα (PDB: 4JPS), iSH2 and nSH2 from PI3Kα (PDB: 3HIZ), and cSH2 from PI3Kβ (PDB: 2Y3A). **C**. Net deuteron differences for all p110α peptides in apo vs pY-activated states for p110α/p85α, p110α/p55α, and p110α/p50α. Significance thresholds as in Fig. 2. Each point represents one peptide. Error bars represent summed S.D. (n = 3 for each). **D**. Net deuteron differences for all p85 peptides spanning the shared region (307-742) in apo vs pY-activated states for the same complexes as in (C). Significance thresholds as in Fig. 2. Each point represents one peptide. Error bars represent summed S.D. (n = 3 for each)

Upon pY activation, increased exchange was observed in regions previously identified during pY activation in the catalytic subunit (Burke et al., 2012), including the helical domain (532–551), ABD–RBD linker (100–119), and C2 domain (456–474) (Fig. 3B,C). An additional increase in exchange was observed in the RBD (223–233) and decrease in exchange in the kinase domain (735-744) in the p55α complex, but not in p85α or p50α. However, these changes were not statistically significant when compared against the pY activated states (i.e. comparing the p110α/p85α pY to p110α/p55α pY), suggesting no major difference in pY activation between isoforms.

In the regulatory subunit, decreased exchange was observed in peptides corresponding to the nSH2 (residues 372–380, 381–400, 414–420) and cSH2 domains (residues 614–637, 647–657, 662–687, 697–710) (Fig. 3B,D). This protection pattern matches the established mechanism of pY peptide activation, where the SH2 domains bind to pYXXM motifs. A single peptide in the nSH2 domain (372–380) showed increased exchange in p85α and p55α but not p50α, consistent with breaking the nSH2-helical interface. However, comparing the pY-bound state of p85α vs p50α to each other did not show a statistically significant change. Overall, these results indicate that PI3Kα responds to pY activation through a conserved mechanism, with no major, global differences attributable to the associated regulatory subunit.

## Discussion

Due to the pivotal role of class IA PI3Ks in human disease, it is essential to understand the molecular details of how a single catalytic subunit can be regulated by all possible class IA regulatory subunits. This has important clinical implications, as multiple allosteric and ATP-competitive inhibitors (André et al., 2019; Buckbinder et al., 2023; Varkaris et al., 2024) are in clinical trials or FDA approved. Understanding differences in regulatory subunit binding is therefore directly relevant to therapeutic development. Our findings show that the lipid kinase activity, membrane-binding affinity, and overall conformational dynamics of p110α are essentially indistinguishable when bound to any of the five class IA PI3K regulatory subunits. Despite differences in domain composition and sequence among p85α, p55α, p50α, p85β, and p55γ, the catalytic subunit exhibits similar behaviour across all complexes. This result underscores that the main regulatory function of the class IA regulatory subunit is primarily driven by the nSH2, iSH2, and cSH2 domains, and suggests that the fundamental mechanism of control is conserved across all regulatory isoforms.

These results reinforce extensive studies on the canonical models of PI3K regulation, which emphasize the importance of the iSH2 and nSH2 domains in stabilizing p110α and maintaining basal inhibition through the nSH2-helical interface, iSH2-C2, and ABD-kinase interfaces (Burke et al., 2012, 2011; Burke and Williams, 2013; Huang et al., 2007; Jenkins et al., 2023; Mandelker et al., 2009; Miled et al., 2007; Torosyan et al., 2025). While most of these studies have been performed in the context of the p85α isoform, our results suggest that these inhibitory interfaces, and mechanism of activation being strongly conserved in p85β and p55γ. Because these inhibitory interactions are retained across all regulatory isoforms, the equal lipid kinase activity is consistent with the HDX-MS results showing similar dynamics of the accessory domains. The additional N-terminal domains present in the longer p85α and p85β isoforms, including the SH3, and BH domains, do not appear to influence p110α’s intrinsic activity or conformational state both in the apo state or upon pY activation. However, it is very important to note that these domains could play important roles in PI3K regulation, specifically interaction with specific polyproline motifs, and in the regulation of p85 complexes that form in the absence of p110 catalytic subunits (Cheung et al., 2011; LoPiccolo et al., 2015).

The absence of detectable differences *in vitro* raises the important question of why multiple regulatory subunits exist. A plausible explanation is that their functional diversification lies outside of direct catalytic regulation of p110α. Instead, isoform-specific contributions may arise from differential expression across tissues, distinct subcellular localization, or scaffolding roles in multi-protein signaling complexes (Kok et al., 2009; Rathinaswamy and Burke, 2019). Multiple isoform-specific post translational modifications have been identified, which provide a mechanism for isoform-specific regulation driven by unique targeting by upstream kinases. This include PKC phosphorylation of p85α in both the nSH2 and cSH2 (Lee et al., 2011), IκB kinase phosphorylation of p85α at S690 (Comb et al., 2012), and Src family kinase phosphorylation of the cSH2 (Cuevas et al., 2001; von Willebrand et al., 1998). Regulatory subunits can also be uniquely targeted for degradation, as the p85β isoform appears to be uniquely degraded downstream of some p110α-selective inhibitors (Song et al., 2022). From this perspective, the regulatory subunits act as interchangeable stabilizers of p110α’s catalytic potential, while their unique ability to function in cellular contexts can be driven by unique isoform-selective interactions with PI3K modulators.

This interpretation has several implications for understanding PI3K signaling in disease. Mutations in p110α that disrupt inhibitory contacts, such as those found in cancer and overgrowth syndromes (Jenkins et al., 2023; Lindhurst et al., 2012; Samuels et al., 2004), would be expected to have similar activating effects with all regulatory subunits. Isoform-specific associations with disease, such as the oncogenic role of p85β (Kim et al., 2023; Liu et al., 2022; Matsubayashi et al., 2024), are therefore likely mediated through mechanisms independent of direct catalytic modulation of the catalytic subunit. Instead, differences may reflect how individual regulatory isoforms organize signaling complexes, recruit effectors, or respond to upstream activators.

Taken together, our results suggest that the diversity of class IA regulatory subunits does not confer differential biochemical control of p110α itself. The evolutionary conservation of the inhibitory mechanism emphasizes the importance of maintaining strict control over PI3K activity in all isoforms, with differences likely driven by additional factors that can specifically target unique regulatory subunits. Future work will be needed to test whether similar equivalence applies to other catalytic isoforms such as p110β and p110δ, and to define the unique isoform-specific regulatory subunit interactions that shape PI3K signaling *in vivo*.

## Acknowledgements

J.E.B. is supported by the Cancer Research Society (Operating grant CRS-1459101). I.B.B is supported by a Natural Sciences and Engineering Research Council of Canada (NSERC) CGS-D scholarship.

## Author Contributions

All biochemical experiments, protein purification, and molecular biology were carried out by I.B.B. HDX-MS experiments were carried out by I.B.B., E.E.W., and H.G.N. Data were analyzed by I.B.B., E.E.W., H.G.N., and J.E.B. Experiments were conceived and designed by J.E.B. and I.B.B. The manuscript was written by J.E.B. and I.B.B. with input from all authors.

## Materials and Methods

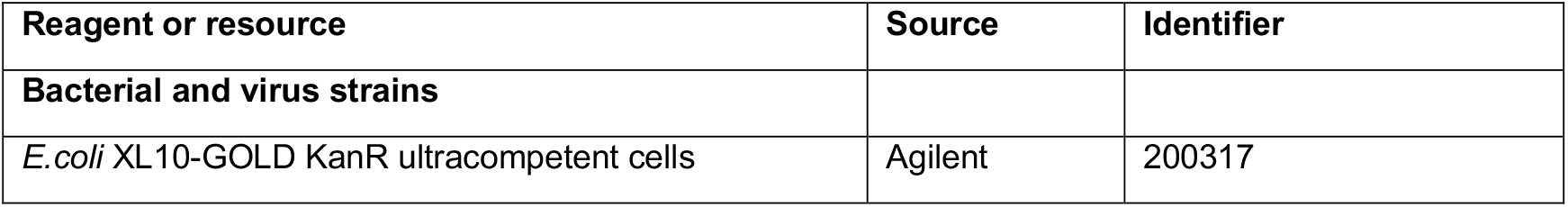
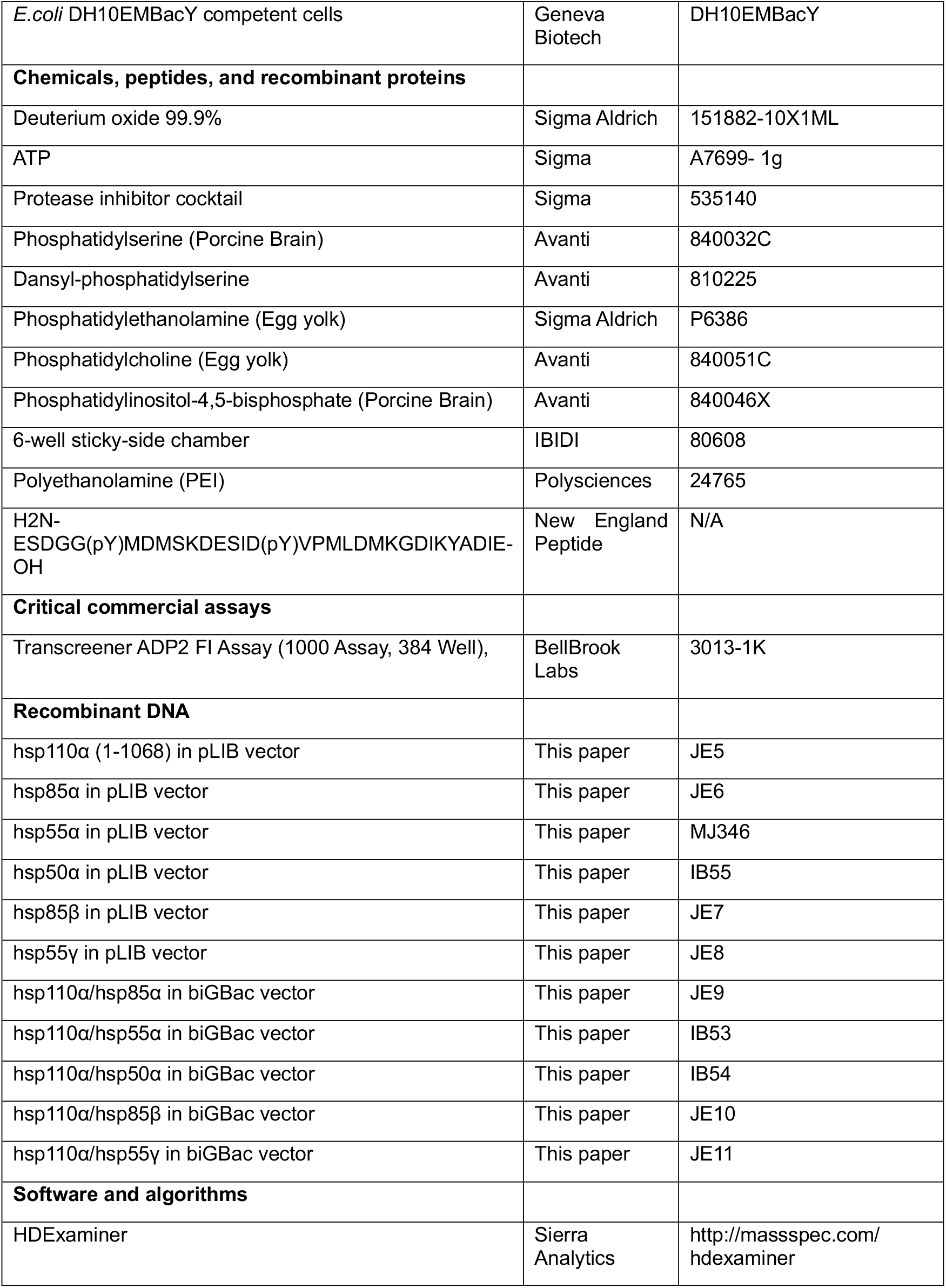
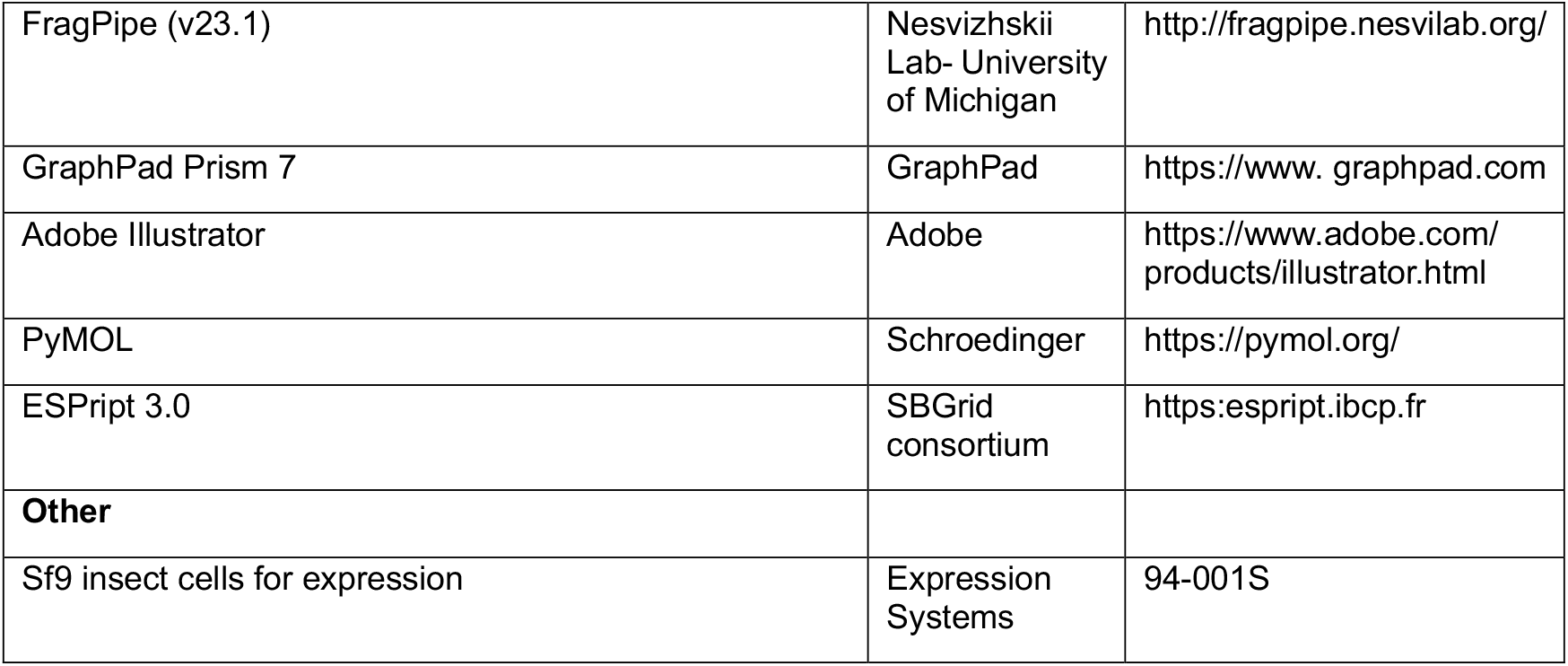

### Plasmid Generation

Genes of interest were inserted into the pLIB vector to allow baculovirus expression in *Spodoptera frugiperda* (Sf9) cells. The full list of all plasmids and reagents used in this manuscript are shown in Table 1. The plasmid containing p110α also expressed a 10X histidine tag N-terminal to the protein, followed by a 2X Strep tag, followed by a Tobacco Etch Virus protease cleavage site. PCR reactions were performed on the WT p85α (*PIK3R1*) and p85β (*PIK3R2*) genes, and PCR purified (Q5 High-Fidelity 2X MasterMix, New England Biosciences #M0492L; QiaQuick PCR Purification Kit, Qiagen #28104). PCR reactions performed on p55γ (*PIK3R3*) used a benign, annotated variant (D338A), and were PCR purified as stated above. Genes were subsequently amplified following the biGBac protocol to generate plasmids containing p110a and a regulatory subunit (p110α/p85α, p110α/p55α, p110α/p50α, p110α/p85β, p110α/p55γ). Single colonies were grown overnight and purified using GeneJET Plasmid Miniprep Kit (Thermo Scientific, #K0503). Plasmid identity was confirmed by sanger sequencing (Plasmidsaurus).

### Virus generation and amplification

The plasmids encoding genes for insect cell expression were transformed into DH10MultiBacY cells (MultiBac, Geneva Biotech) to generate bacmid containing the genes of interest. Successful generation was identified by blue-white colony screening and the bacmid was purified using a standard isopropanol-ethanol extraction method.

Briefly, colonies were grown overnight (∼16 h) in 3–5 mL 2xYT (BioBasic #SD7019). Cells were pelleted by centrifugation and the pellet was resuspended in 225 μL Resuspension Solution (Thermo Scientific, #K0503), chemically lysed by the addition of 225 μL Lysis Solution, and the lysis reaction was neutralised by addition of 300 μL Neutralization Solution. Following centrifugation at 21130 rcf and 4 °C (Rotor #5424 R), the supernatant was separated and mixed with 600 μL isopropanol to precipitate the DNA out of solution. Further centrifugation at the same temperature and speed pelleted the bacmid DNA, which was then washed with 500 μL 70% ethanol three times. The Bacmid DNA pellet was then dried for 1 min and re-suspended in 50 μL Elution Buffer.

Purified bacmid was then transfected into Sf9 cells. 2 mL of Sf9 cells at 1.0 × 10^6^ cells/mL were aliquoted into the wells of a six-well plate and allowed to attach, creating a monolayer of cells at ∼70–80% confluency. Transfection reactions were prepared by the addition of 2–10 μg of bacmid DNA to 200 μl 1xPBS and 7 μL polyethyleneimine (PEI) at 1 mg/mL (Polyethyleneimine “Max” MW 40,000, Polysciences #24765, USA). solution was mixed, and the reaction occurred for 20–30 min before addition drop-by-drop to an Sf9 monolayer containing well. Transfections were allowed to proceed for 4–5 days before harvesting virus containing supernatant as a P1 viral stock.

Viral stocks were amplified by adding P1 viral stock to suspension Sf9 cells between 1– 2 × 10^6^ cells/mL at a 1/100 volume ratio. This amplification produces a P2 stage viral stock that can be used in final protein expression. The amplification proceeded for 3–4 days before harvesting, with cell shaking at 120 RPM in a 27 °C shaker (New Brunswick). Harvesting of P2 viral stocks was carried out by centrifuging cell suspensions in 50 mL Falcon tubes at 2281 RCF (Beckman GS-15), collecting the supernatant in a fresh sterile tube, and adding 5–10% inactivated foetal bovine serum (FBS; VWR Canada #97068-085).

### Expression and purification of recombinant proteins

All PI3Kα constructs were purified by expressing the catalytic subunit and the regulatory subunit together using the biGBac expression system in Sf9 cells. After expressing the cells at 27 °C for 55 hours, the cells were harvested at 1739 × *g* at 4 °C using Eppendorf Centrifuge 5810R and the cells were flash frozen using liquid nitrogen and stored in −80 °C.

Frozen insect cell pellets were resuspended in lysis buffer (20 mM Tris pH 8.0, 100 mM NaCl, 10 mM imidazole pH 8.0, 5% glycerol (v/v), 2 mM βME), protease inhibitor (Protease Inhibitor Cocktail Set III, Sigma)) and sonicated for 3 minutes (15s on, 15s off, level 4.0, Misonix sonicator 3000). Triton-X was added to the lysate to a final concentration of 0.1% and clarified by spinning at 15,000 RCF at 4°C for 45 minutes (Beckman Coulter JA-20 rotor). The supernatant was loaded onto a 5 mL HisTrap™ FF crude column (GE Healthcare) equilibrated in NiNTA A buffer (20 mM Tris pH 8.0, 100 mM NaCl, 20 mM imidazole pH 8.0, 5% (v/v) glycerol, 2 mM βME). The column was washed with high salt NiNTA A buffer (20 mM Tris pH 8.0, 1 M NaCl, 20 mM imidazole pH 8.0, 5% (v/v) glycerol, 2 mM βME), NiNTA A buffer, 6% NiNTA B buffer (20 mM Tris pH 8.0, 100 mM NaCl, 250 mM imidazole pH 8.0, 5% (v/v) glycerol, 2 mM βME) and the protein was eluted with 100% NiNTA B. The eluent was loaded onto a 5 mL StrepTrap™ HP column (GE Healthcare) equilibrated in gel filtration buffer (20 mM HEPES 7.5, 150 mM NaCl, 5% glycerol, 0.5 mM TCEP). The column was washed with gel filtration buffer and loaded with tobacco etch virus protease. After cleavage on the column for three hours, the PI3Kα protein constructs were eluted in gel filtration buffer. The protein was concentrated in a 50,000 MWCO Amicon Concentrator (Millipore) to <1 mL and injected onto a Superdex™ 200 10/300 GL Increase size-exclusion column (GE Healthcare) equilibrated in gel filtration buffer. After size exclusion, the protein was concentrated, aliquoted, frozen, and stored at −80°C.

### Lipid vesicle preparation

Three sets of vesicles were prepared for kinase assays, protein-lipid FRET assays, and HDX-MS: 5% brain phosphatidylinositol 4,5-bisphosphate (PIP_2_) and 95% porcine brain phosphatidylserine (PS) for kinase assays; 5% porcine brain PIP_2_, 25% porcine brain PS, 60% egg yolk PE, and 10% dansyl PS for protein-lipid FRET assays; and 5% PIP_2_, 30% brain PS, 50% egg yolk phosphatidylethanolamine (PE), and 15% egg yolk phosphatidylcholine (PC) for HDX-MS. To generate vesicles, the lipid mixtures were combined in organic solvent. The mixture was then evaporated using a stream of argon gas followed by desiccation under vacuum for 45 min. The lipids were resuspended in lipid buffer (20 mM HEPES 7.5, 100 mM KCl, 0.5 mM EDTA) and the solution was vortexed for 5 minutes followed by sonication for 15 minutes. The vesicles were then subjected to ten freeze thaw cycles and extruded 11 times through a 100-nm filter (Avanti Research: 610000-1EA). The extruded vesicles were sub-aliquoted and stored at −80°C. Final vesicle concentration was 5 mg/mL.

### Lipid kinase assays

All ATPase assays used the Transcreener ADP2 Fluorescence Intensity (FI) assay (Bellbrook labs) which measures formation of ADP. In all, 2 μL of a 2x PI3Kα kinase solution (final concentration 5 nM) was mixed with 2 μL of a 2x substrate solution containing ATP (final concentration 100 µM) and PIP2 vesicles (final concentration 0.45 mg/mL), and the reaction was allowed to proceed for 60 min at 20 °C. After the 60-minute incubation, all reactions were stopped with 4 μL of 2X stop and detect solution containing Stop and Detect buffer (20mM HEPES, 0.02% Brij-35, 400mM 40mM EDTA pH 7.5), 8 nM ADP Alexa Fluor 594 Tracer and 93.7 μg/mL ADP2 Antibody IRDye QC-1), covered and incubated at 20°C for 60 min before reading the fluorescence. The fluorescence intensity was measured using an Agilent BioTek Synergy H1 plate reader at excitation 585/20 nm and emission 626/20 nm. This data was normalised against a 0–100% ADP window made using conditions containing a final concentration of 100 μM ATP or ADP. % ATP turnover was interpolated from an ATP standard curve obtained from performing the assay on 100 μM (total) ATP/ADP mixtures with increasing concentrations of ADP.

### Protein-lipid FRET assays

Protein-lipid FRET experiments were carried out at 250 nM PI3Kα. Assays were initiated by mixing 2.5 µL PI3Ka (final concentration of 250 nM) with 2.5 µL of pY peptide diluted in protein buffer [20 mM HEPES pH 7.5, 150 mM NaCl, 5% glycerol [v/v], 0.5 mM TCEP] (final concentration of 1 µM) or protein buffer for 15 mins at 20 °C. 5 µL of protein-lipid FRET vesicles (PIP_2_/PE/PS/dansyl PS, final concentration of 16.67 µg/mL) diluted in lipid buffer were added to the protein-pY mixture and were incubated for 15 mins at 20 °C. The plate was then read using a SpectraMax M5 plate reader using a 280-nm excitation filter with 350 nm and 520 nm emission filters to measure Trp and Dansyl-PS FRET emissions, respectively. The FRET signal shown in Fig. 1G has normalized I-Io along the Y axis where I is the intensity of 520 with protein and Io is the intensity of lipid alone.

### HDX-MS analysis: sample preparation

HDX-MS reactions comparing p110α/p85α, p110α/p85β, and p110α/p55γ were conducted in 20 μl reaction volumes with a final PI3K amount of 10 pmol. Prior to HD exchange, 4 μl of each protein was mixed with 1 uL of 75 μM pY peptide or pY buffer and 1 uL of 15% DMSO in GFB and allowed to incubate for 15 minutes on ice. To initiate HD exchange, 14 μl of D_2_O buffer [20 µL 1.0 M HEPES pH 7.5 [20 mM], 30 µL 5 M NaCl [150 mM], 1 uL 0.5 M TCEP [0.5 mM], 1 mL D2O [92.68%], 26.9 uL 100% DMSO for final 2.5% DMSO] was added to the protein mix (final D_2_O concentration of 64.87%). Exchange was carried out for 3, 300 and 3000 s at 20 °C.

HDX reactions comparing full-length p110α/p85α against p110α/p55α and p110α/p50α with were conducted in 20 μl reaction volumes with a final PI3K amount of 15 pmol. Prior to HD exchange, 3 μl of protein was incubated with 1 μl of 50 μM pY peptide or pY buffer and 2 uL of 5% PIP_2_/30% PS/50% PE/15% PC membrane at 5 mg/mL or lipid buffer and allowed to incubate for 10 min at 20°C. Exchange was initiated by the addition of 14 μl of D_2_O buffer to the protein + /− pY mixture (final D2O concentration of 64.87%). Exchange was carried out for 3, 30, and 300 at 20 °C.

All conditions and timepoints were generated in independent triplicate. All exchange reactions were terminated by the addition of ice-cold quench buffer to give a final concentration 0.6 M guanidine-HCl and 0.9% formic acid. Samples were flash frozen in liquid nitrogen immediately after quenching and stored at −80 °C until injected onto the ultra-performance liquid chromatography (UPLC) system for proteolytic cleavage, peptide separation, and injection onto a QTOF for mass analysis, described below.

### HDX-MS analysis: protein digestion and tandem MS data collection

Protein samples were rapidly thawed and injected onto an integrated fluidics system containing an HDx-3 PAL liquid handling robot and climate-controlled (2 °C) chromatography system (LEAP Technologies), a Dionex Ultimate 3000 UHPLC system, and an Impact HD QTOF or an Impact II QTOF mass spectrometer (Bruker). The full details of the automated LC system are described in (Stariha et al., 2021). The protein was run over two immobilised pepsin columns (Affipro; AP-PC-001) at 200 μl/min for 4 min at 2°C. The resulting peptides were collected and desalted on a C18 trap column (ACQUITY UPLC BEH C18 1.7 μm column, 2.1 mm × 5 mm; Waters 186004629). The trap was subsequently eluted in line with an ACQUITY 1.7 μm particle, 2.1 mm × 100 mm C18 UPLC column (Waters; 186003686), using a gradient of 3–35% B (Buffer A 0.1% formic acid; Buffer B 100% acetonitrile) over 11 min immediately followed by a gradient of 35–80% over 5 min. MS experiments acquired over a mass range from 150 to 2200 mass/charge ratio (m/z) using an electrospray ionisation source operated at a temperature of 200 °C and a spray voltage of 4.5 kV.

### HDX-MS analysis: peptide identification

Peptides were identified from the non-deuterated samples of p110α/p85α complex for WT and other mutants using data-dependent acquisition following tandem MS (MS/MS) experiments (0.5-s precursor scan from 150 to 2000 m/z: 12 0.25 s fragment scans from 150 to 2000 m/z). MS/MS datasets were analysed using FragPipe (v23.1), and peptide identification was carried out by using a false discovery–based approach, with a threshold set to 1% using a database of purified proteins and known contaminants found in SF9 cells (Dobbs et al., 2020; Kong et al., 2017). MSFragger was used, and the precursor mass tolerance error was set to −20 to 20 ppm. The fragment mass tolerance was set at 20 ppm. Protein digestion was set as nonspecific, searching between lengths of 4 and 50 aa, with a mass range of 400–5000 Da.

### HDX-MS analysis: mass analysis of peptide centroids and measurement of deuterium incorporation

HD-Examiner Software (Sierra Analytics) was used to automatically calculate the level of deuterium incorporation into each peptide. All peptides were manually inspected for correct charge state, correct retention time, and appropriate selection of isotopic distribution. Deuteration levels were calculated using the centroid of the experimental isotope clusters. HDX-MS results are presented with no correction for back exchange shown in the Source data, with the only correction being applied correcting for the deuterium oxide percentage of the buffer used in the exchange (64.9%).

Differences in exchange in a peptide were considered significant if they met all three of the following criteria for apo and pY conditions: ≥5% change in exchange, ≥0.45 Da difference in exchange, and a *P*-value of <0.01 using a two-tailed Student’s *t* test. The raw data for all analysed peptides is available in the source data.

The differences in deuterium exchange are visualised in different ways. To allow for visualisation of differences across conditions, we used number of deuteron difference (#D) plots (Figs. 2A-C + 3C-D). These plots show the total difference in deuterium incorporation over the entire HDX time course, with each point indicating a single peptide. These graphs are calculated by summing the differences at every time point for each peptide and propagating the error. For a selection of peptides, we are showing the %D incorporation over a time course, which allows for comparison of multiple conditions at the same time for a given region (Fig. 2D). Samples were only compared when they were set at the same time and were never compared to experiments completed with a different final D_2_O level. The data analysis statistics for all HDX-MS experiments are in the source data according to published guidelines (Masson et al., 2019). The HDX-MS proteomics data generated in this study have been deposited to the ProteomeXchange Consortium via the PRIDE partner repository (Perez-Riverol et al., 2019).

## References

André, F., Ciruelos, E., Rubovszky, G., Campone, M., Loibl, S., Rugo, H.S., Iwata, H., Conte, P., Mayer, I.A., Kaufman, B., Yamashita, T., Lu, Y.-S., Inoue, K., Takahashi, M., Pápai, Z., Longin, A.-S., Mills, D., Wilke, C., Hirawat, S., Juric, D., SOLAR-1 Study Group, 2019. Alpelisib for PIK3CA-Mutated, Hormone Receptor-Positive Advanced Breast Cancer. N. Engl. J. Med. 380, 1929–1940. 10.1056/NEJMoa1813904

Buckbinder, L., St Jean, D.J., Tieu, T., Ladd, B., Hilbert, B., Wang, W., Alltucker, J.T., Manimala, S., Kryukov, G.V., Brooijmans, N., Dowdell, G., Jonsson, P., Huff, M., Guzman-Perez, A., Jackson, E.L., Goncalves, M.D., Stuart, D.D., 2023. STX-478, a Mutant-Selective, Allosteric PI3Kα Inhibitor Spares Metabolic Dysfunction and Improves Therapeutic Response in PI3Kα-Mutant Xenografts. Cancer Discov 13, 2432–2447. 10.1158/2159-8290.CD-23-0396

Buckles, T.C., Ziemba, B.P., Masson, G.R., Williams, R.L., Falke, J.J., 2017. Single-Molecule Study Reveals How Receptor and Ras Synergistically Activate PI3Kα and PIP3 Signaling. Biophys J 113, 2396–2405. 10.1016/j.bpj.2017.09.018

Burke, J.E., 2018. Structural Basis for Regulation of Phosphoinositide Kinases and Their Involvement in Human Disease. Molecular Cell 71, 653–673. 10.1016/j.molcel.2018.08.005

Burke, J.E., Perisic, O., Masson, G.R., Vadas, O., Williams, R.L., 2012. Oncogenic mutations mimic and enhance dynamic events in the natural activation of phosphoinositide 3-kinase p110α (PIK3CA). Proc. Natl. Acad. Sci. U.S.A. 109, 15259–15264. 10.1073/pnas.1205508109

Burke, J.E., Vadas, O., Berndt, A., Finegan, T., Perisic, O., Williams, R.L., 2011. Dynamics of the phosphoinositide 3-kinase p110δ interaction with p85α and membranes reveals aspects of regulation distinct from p110α. Structure 19, 1127–1137. 10.1016/j.str.2011.06.003

Burke, J.E., Williams, R.L., 2015. Synergy in activating class I PI3Ks. Trends Biochem Sci 40, 88–100. 10.1016/j.tibs.2014.12.003

Burke, J.E., Williams, R.L., 2013. Dynamic steps in receptor tyrosine kinase mediated activation of class IA phosphoinositide 3-kinases (PI3K) captured by H/D exchange (HDX-MS). Adv Biol Regul 53, 97–110. 10.1016/j.jbior.2012.09.005

Cheung, L., Hennessy, B., Li, J., Yu, S., Myers, A., Djordjevic, B., Lu, Y., Stemke-Hale, K., Zhang, F., Ju, Z., Cantley, L., Scherer, S., Liang, H., Lu, K., Broaddus, R., Mills, G., 2011. High Frequency of PIK3R1 and PIK3R2 Mutations in Endometrial Cancer Elucidates a Novel Mechanism for Regulation of PTEN Protein Stability. Cancer Discov 1, 170–185. 10.1158/2159-8290.CD-11-0039

Comb, W.C., Hutti, J.E., Cogswell, P., Cantley, L.C., Baldwin, A.S., 2012. p85α SH2 domain phosphorylation by IKK promotes feedback inhibition of PI3K and Akt in response to cellular starvation. Mol. Cell 45, 719–730. 10.1016/j.molcel.2012.01.010

Cortés, I., Sánchez-Ruíz, J., Zuluaga, S., Calvanese, V., Marqués, M., Hernández, C., Rivera, T., Kremer, L., González-García, A., Carrera, A.C., 2012. p85β phosphoinositide 3-kinase subunit regulates tumor progression. Proc Natl Acad Sci U S A 109, 11318– 11323. 10.1073/pnas.1118138109

Cuevas, B.D., Lu, Y., Mao, M., Zhang, J., LaPushin, R., Siminovitch, K., Mills, G.B., 2001. Tyrosine phosphorylation of p85 relieves its inhibitory activity on phosphatidylinositol 3-kinase. J. Biol. Chem. 276, 27455–27461. 10.1074/jbc.M100556200

Dobbs, J.M., Jenkins, M.L., Burke, J.E., 2020. Escherichia coli and Sf9 Contaminant Databases to Increase Efficiency of Tandem Mass Spectrometry Peptide Identification in Structural Mass Spectrometry Experiments. J. Am. Soc. Mass Spectrom. 31, 2202–2209. 10.1021/jasms.0c00283

Dornan, G.L., Siempelkamp, B.D., Jenkins, M.L., Vadas, O., Lucas, C.L., Burke, J.E., 2017. Conformational disruption of PI3Kδ regulation by immunodeficiency mutations in PIK3CD and PIK3R1. Proc. Natl. Acad. Sci. U.S.A. 114, 1982–1987. 10.1073/pnas.1617244114

Dornan, G.L., Stariha, J.T.B., Rathinaswamy, M.K., Powell, C.J., Boulanger, M.J., Burke, J.E., 2020. Defining How Oncogenic and Developmental Mutations of PIK3R1 Alter the Regulation of Class IA Phosphoinositide 3-Kinases. Structure 28, 145-156.e5. 10.1016/j.str.2019.11.013

Fox, M., Mott, H.R., Owen, D., 2020. Class IA PI3K regulatory subunits: p110-independent roles and structures. Biochem Soc Trans 48, 1397–1417. 10.1042/BST20190845

Gong, G.Q., Bilanges, B., Allsop, B., Masson, G.R., Roberton, V., Askwith, T., Oxenford, S., Madsen, R.R., Conduit, S.E., Bellini, D., Fitzek, M., Collier, M., Najam, O., He, Z., Wahab, B., McLaughlin, S.H., Chan, A.W.E., Feierberg, I., Madin, A., Morelli, D., Bhamra, A., Vinciauskaite, V., Anderson, K.E., Surinova, S., Pinotsis, N., Lopez-Guadamillas, E., Wilcox, M., Hooper, A., Patel, C., Whitehead, M.A., Bunney, T.D., Stephens, L.R., Hawkins, P.T., Katan, M., Yellon, D.M., Davidson, S.M., Smith, D.M., Phillips, J.B., Angell, R., Williams, R.L., Vanhaesebroeck, B., 2023. A small-molecule PI3Kα activator for cardioprotection and neuroregeneration. Nature 618, 159–168. 10.1038/s41586-023-05972-2

Harris, N.J., Jenkins, M.L., Nam, S.-E., Rathinaswamy, M.K., Parson, M.A.H., Ranga-Prasad, H., Dalwadi, U., Moeller, B.E., Sheeky, E., Hansen, S.D., Yip, C.K., Burke, J.E., 2023. Allosteric activation or inhibition of PI3Kγ mediated through conformational changes in the p110γ helical domain. Elife 12, RP88058. 10.7554/eLife.88058

Huang, C., Mandelker, D., Schmidt-Kittler, O., Samuels, Y., Velculescu, V., Kinzler, K., Vogelstein, B., Gabelli, S., Amzel, L., 2007. The structure of a human p110alpha/p85alpha complex elucidates the effects of oncogenic PI3Kalpha mutations. Science 318, 1744–1748. 10.1126/science.1150799

Inukai, K., Funaki, M., Ogihara, T., Katagiri, H., Kanda, A., Anai, M., Fukushima, Y., Hosaka, T., Suzuki, M., Shin, B.C., Takata, K., Yazaki, Y., Kikuchi, M., Oka, Y., Asano, T., 1997. p85alpha gene generates three isoforms of regulatory subunit for phosphatidylinositol 3-kinase (PI 3-Kinase), p50alpha, p55alpha, and p85alpha, with different PI 3-kinase activity elevating responses to insulin. J Biol Chem 272, 7873–7882. 10.1074/jbc.272.12.7873

Jenkins, M.L., Ranga-Prasad, H., Parson, M.A.H., Harris, N.J., Rathinaswamy, M.K., Burke, J.E., 2023. Oncogenic mutations of PIK3CA lead to increased membrane recruitment driven by reorientation of the ABD, p85 and C-terminus. Nat Commun 14, 181. 10.1038/s41467-023-35789-6

Kim, C.-W., Lee, J.M., Park, S.W., 2023. Divergent roles of the regulatory subunits of class IA PI3K. Front Endocrinol (Lausanne) 14, 1152579. 10.3389/fendo.2023.1152579

Kok, K., Geering, B., Vanhaesebroeck, B., 2009. Regulation of phosphoinositide 3-kinase expression in health and disease. Trends in Biochemical Sciences 34, 115–127. 10.1016/j.tibs.2009.01.003

Kong, A.T., Leprevost, F.V., Avtonomov, D.M., Mellacheruvu, D., Nesvizhskii, A.I., 2017. MSFragger: ultrafast and comprehensive peptide identification in mass spectrometry–based proteomics. Nat Methods 14, 513–520. 10.1038/nmeth.4256

Lee, J., Chiu, Y., Asara, J., Cantley, L., 2011. Inhibition of PI3K binding to activators by serine phosphorylation of PI3K regulatory subunit p85{alpha} Src homology-2 domains. Proc. Natl. Acad. Sci. U.S.A. 108, 14157–14162. 10.1073/pnas.1107747108/-/DCSupplemental

Lindhurst, M.J., Parker, V.E.R., Payne, F., Sapp, J.C., Rudge, S., Harris, J., Witkowski, A.M., Zhang, Q., Groeneveld, M.P., Scott, C.E., Daly, A., Huson, S.M., Tosi, L.L., Cunningham, M.L., Darling, T.N., Geer, J., Gucev, Z., Sutton, V.R., Tziotzios, C., Dixon, A.K., Helliwell, T., O’Rahilly, S., Savage, D.B., Wakelam, M.J.O., Barroso, I., Biesecker, L.G., Semple, R.K., 2012. Mosaic overgrowth with fibroadipose hyperplasia is caused by somatic activating mutations in PIK3CA. Nat. Genet. 44, 928–933. 10.1038/ng.2332

Liu, Y., Wang, D., Li, Z., Li, X., Jin, M., Jia, N., Cui, X., Hu, G., Tang, T., Yu, Q., 2022. Pan-cancer analysis on the role of PIK3R1 and PIK3R2 in human tumors. Sci Rep 12, 5924. 10.1038/s41598-022-09889-0

LoPiccolo, J., Kim, S.J., Shi, Y., Wu, B., Wu, H., Chait, B.T., Singer, R.H., Sali, A., Brenowitz, M., Bresnick, A.R., Backer, J.M., 2015. Assembly and Molecular Architecture of the Phosphoinositide 3-Kinase p85α Homodimer. J Biol Chem 290, 30390–30405. 10.1074/jbc.M115.689604

Madsen, R.R., Vanhaesebroeck, B., 2020. Cracking the context-specific PI3K signaling code. Science Signaling 13, eaay2940. 10.1126/scisignal.aay2940

Mandelker, D., Gabelli, S.B., Schmidt-Kittler, O., Zhu, J., Cheong, I., Huang, C.-H., Kinzler, K.W., Vogelstein, B., Amzel, L.M., 2009. A frequent kinase domain mutation that changes the interaction between PI3Kalpha and the membrane. Proc Natl Acad Sci U S A 106, 16996–17001. 10.1073/pnas.0908444106

Manning, B.D., Toker, A., 2017. AKT/PKB Signaling: Navigating the Network. Cell 169, 381– 405. 10.1016/j.cell.2017.04.001

Masson, G.R., Burke, J.E., Ahn, N.G., Anand, G.S., Borchers, C., Brier, S., Bou-Assaf, G.M., Engen, J.R., Englander, S.W., Faber, J., Garlish, R., Griffin, P.R., Gross, M.L., Guttman, M., Hamuro, Y., Heck, A.J.R., Houde, D., Iacob, R.E., Jørgensen, T.J.D., Kaltashov, I.A., Klinman, J.P., Konermann, L., Man, P., Mayne, L., Pascal, B.D., Reichmann, D., Skehel, M., Snijder, J., Strutzenberg, T.S., Underbakke, E.S., Wagner, C., Wales, T.E., Walters, B.T., Weis, D.D., Wilson, D.J., Wintrode, P.L., Zhang, Z., Zheng, J., Schriemer, D.C., Rand, K.D., 2019. Recommendations for performing, interpreting and reporting hydrogen deuterium exchange mass spectrometry (HDX-MS) experiments. Nat Methods 16, 595–602. 10.1038/s41592-019-0459-y

Masson, G.R., Jenkins, M.L., Burke, J.E., 2017. An overview of hydrogen deuterium exchange mass spectrometry (HDX-MS) in drug discovery. Expert Opin Drug Discov 12, 981–994. 10.1080/17460441.2017.1363734

Matsubayashi, H.T., Mountain, J., Takahashi, N., Deb Roy, A., Yao, T., Peterson, A.F., Saez Gonzalez, C., Kawamata, I., Inoue, T., 2024. Non-catalytic role of phosphoinositide 3-kinase in mesenchymal cell migration through non-canonical induction of p85β/AP2-mediated endocytosis. Nat Commun 15, 2612. 10.1038/s41467-024-46855-y

Miled, N., Yan, Y., Hon, W.-C., Perisic, O., Zvelebil, M., Inbar, Y., Schneidman-Duhovny, D., Wolfson, H.J., Backer, J.M., Williams, R.L., 2007. Mechanism of two classes of cancer mutations in the phosphoinositide 3-kinase catalytic subunit. Science 317, 239–242. 10.1126/science.1135394

Nicolau-Neto, P., Da Costa, N.M., de Souza Santos, P.T., Gonzaga, I.M., Ferreira, M.A., Guaraldi, S., Moreira, M.A., Seuánez, H.N., Brewer, L., Bergmann, A., Boroni, M., Mencalha, A.L., Kruel, C.D.P., Lima, S.C.S., Esposito, D., Simão, T.A., Pinto, L.F.R., 2018. Esophageal squamous cell carcinoma transcriptome reveals the effect of FOXM1 on patient outcome through novel PIK3R3 mediated activation of PI3K signaling pathway. Oncotarget 9, 16634–16647. 10.18632/oncotarget.24621

Peng, Y.-P., Zhu, Y., Yin, L.-D., Wei, J.-S., Liu, X.-C., Zhu, X.-L., Miao, Y., 2018. PIK3R3 Promotes Metastasis of Pancreatic Cancer via ZEB1 Induced Epithelial-Mesenchymal Transition. Cellular Physiology and Biochemistry 46, 1930–1938. 10.1159/000489382

Perez-Riverol, Y., Csordas, A., Bai, J., Bernal-Llinares, M., Hewapathirana, S., Kundu, D.J., Inuganti, A., Griss, J., Mayer, G., Eisenacher, M., Pérez, E., Uszkoreit, J., Pfeuffer, J., Sachsenberg, T., Yılmaz, Ş., Tiwary, S., Cox, J., Audain, E., Walzer, M., Jarnuczak, A.F., Ternent, T., Brazma, A., Vizcaíno, J.A., 2019. The PRIDE database and related tools and resources in 2019: improving support for quantification data. Nucleic Acids Res 47, D442–D450. 10.1093/nar/gky1106

Piccione, E., Case, R.D., Domchek, S.M., Hu, P., Chaudhuri, M., Backer, J.M., Schlessinger, J., Shoelson, S.E., 1993. Phosphatidylinositol 3-kinase p85 SH2 domain specificity defined by direct phosphopeptide/SH2 domain binding. Biochemistry 32, 3197– 3202. 10.1021/bi00064a001

Pons, S., Asano, T., Glasheen, E., Miralpeix, M., Zhang, Y., Fisher, T.L., Myers, M.G., Sun, X.J., White, M.F., 1995. The structure and function of p55PIK reveal a new regulatory subunit for phosphatidylinositol 3-kinase. Mol Cell Biol 15, 4453–4465. 10.1128/MCB.15.8.4453

Rathinaswamy, M.K., Burke, J.E., 2019. Class I phosphoinositide 3-kinase (PI3K) regulatory subunits and their roles in signaling and disease. Adv Biol Regul 100657. 10.1016/j.jbior.2019.100657

Rathinaswamy, M.K., Dalwadi, U., Fleming, K.D., Adams, C., Stariha, J.T.B., Pardon, E., Baek, M., Vadas, O., DiMaio, F., Steyaert, J., Hansen, S.D., Yip, C.K., Burke, J.E., 2021a. Structure of the phosphoinositide 3-kinase (PI3K) p110γ-p101 complex reveals molecular mechanism of GPCR activation. Sci Adv 7, eabj4282. 10.1126/sciadv.abj4282

Rathinaswamy, M.K., Fleming, K.D., Dalwadi, U., Pardon, E., Harris, N.J., Yip, C.K., Steyaert, J., Burke, J.E., 2021b. HDX-MS-optimized approach to characterize nanobodies as tools for biochemical and structural studies of class IB phosphoinositide 3-kinases. Structure 29, 1371-1381.e6. 10.1016/j.str.2021.07.002

Rathinaswamy, M.K., Gaieb, Z., Fleming, K.D., Borsari, C., Harris, N.J., Moeller, B.E., Wymann, M.P., Amaro, R.E., Burke, J.E., 2021c. Disease-related mutations in PI3Kγ disrupt regulatory C-terminal dynamics and reveal a path to selective inhibitors. Elife 10, e64691. 10.7554/eLife.64691

Rathinaswamy, M.K., Jenkins, M.L., Duewell, B.R., Zhang, X., Harris, N.J., Evans, J.T., Stariha, J.T.B., Dalwadi, U., Fleming, K.D., Ranga-Prasad, H., Yip, C.K., Williams, R.L., Hansen, S.D., Burke, J.E., 2023. Molecular basis for differential activation of p101 and p84 complexes of PI3Kγ by Ras and GPCRs. Cell Rep 42, 112172. 10.1016/j.celrep.2023.112172

Samuels, Y., Wang, Z., Bardelli, A., Silliman, N., Ptak, J., Szabo, S., Yan, H., Gazdar, A., Powell, S., Riggins, G., Willson, J., Markowitz, S., Kinzler, K., Vogelstein, B., Velculescu, V., 2004. High frequency of mutations of the PIK3CA gene in human cancers. Science 304, 554. 10.1126/science.1096502

Shaw, A.L., Barlow-Busch, I., Burke, J.E., 2025. Molecular basis for regulation of the class I phosphoinositide 3-kinases (PI3Ks), and their targeting in human disease. Biochimica et Biophysica Acta (BBA) - Molecular and Cell Biology of Lipids 1870, 159689. 10.1016/j.bbalip.2025.159689

Shaw, A.L., Burke, J.E., 2025. Molecular insight on the role of the phosphoinositide PIP3 in regulating the protein kinases Akt, PDK1, and BTK. Biochem Soc Trans BST20253059. 10.1042/BST20253059

Siempelkamp, B.D., Rathinaswamy, M.K., Jenkins, M.L., Burke, J.E., 2017. Molecular mechanism of activation of class IA phosphoinositide 3-kinases (PI3Ks) by membrane-localized HRas. J Biol Chem 292, 12256–12266. 10.1074/jbc.M117.789263

Song, K.W., Edgar, K.A., Hanan, E.J., Hafner, M., Oeh, J., Merchant, M., Sampath, D., Nannini, M.A., Hong, R., Phu, L., Forrest, W.F., Stawiski, E., Schmidt, S., Endres, N., Guan, J., Wallin, J.J., Cheong, J., Plise, E.G., Lewis Phillips, G.D., Salphati, L., Heffron, T.P., Olivero, A.G., Malek, S., Staben, S.T., Kirkpatrick, D.S., Dey, A., Friedman, L.S., 2022. RTK-Dependent Inducible Degradation of Mutant PI3Kα Drives GDC-0077 (Inavolisib) Efficacy. Cancer Discov 12, 204–219. 10.1158/2159-8290.CD-21-0072

Stariha, J.T.B., Hoffmann, R.M., Hamelin, D.J., Burke, J.E., 2021. Probing Protein–Membrane Interactions and Dynamics Using Hydrogen–Deuterium Exchange Mass Spectrometry (HDX-MS), in:Daviter, T., Johnson, C.M., McLaughlin, S.H., Williams, M.A. (Eds.), Protein-Ligand Interactions: Methods and Applications, Methods in Molecular Biology. Springer US, New York, NY, pp. 465–485. 10.1007/978-1-0716-1197-5_22

Torosyan, H., Paul, M.D., Maker, A., Meyer, B.G., Jura, N., Verba, K.A., 2025. Structures of the PI3Kα/KRas complex on lipid bilayers reveal the molecular mechanism of PI3Kα activation. bioRxiv 2025.03.22.644753. 10.1101/2025.03.22.644753

Ueki, K., Fruman, D.A., Brachmann, S.M., Tseng, Y.-H., Cantley, L.C., Kahn, C.R., 2002. Molecular balance between the regulatory and catalytic subunits of phosphoinositide 3-kinase regulates cell signaling and survival. Mol Cell Biol 22, 965–977. 10.1128/MCB.22.3.965-977.2002

Vallejo-Díaz, J., Chagoyen, M., Olazabal-Morán, M., González-García, A., Carrera, A.C., 2019. The Opposing Roles of PIK3R1/p85α and PIK3R2/p85β in Cancer. Trends in Cancer 5, 233–244. 10.1016/j.trecan.2019.02.009

Varkaris, A., Pazolli, E., Gunaydin, H., Wang, Q., Pierce, L., Boezio, A.A., Bulku, A., DiPietro, L., Fridrich, C., Frost, A., Giordanetto, F., Hamilton, E.P., Harris, K., Holliday, M., Hunter, T.L., Iskandar, A., Ji, Y., Larivée, A., LaRochelle, J.R., Lescarbeau, A., Llambi, F., Lormil, B., Mader, M.M., Mar, B.G., Martin, I., McLean, T.H., Michelsen, K., Pechersky, Y., Puente-Poushnejad, E., Raynor, K., Rogala, D., Samadani, R., Schram, A.M., Shortsleeves, K., Swaminathan, S., Tajmir, S., Tan, G., Tang, Y., Valverde, R., Wehrenberg, B., Wilbur, J., Williams, B.R., Zeng, H., Zhang, H., Walters, W.P., Wolf, B.B., Shaw, D.E., Bergstrom, D.A., Watters, J., Fraser, J.S., Fortin, P.D., Kipp, D.R., 2024. Discovery and Clinical Proof-of-Concept of RLY-2608, a First-in-Class Mutant-Selective Allosteric PI3Kα Inhibitor That Decouples Antitumor Activity from Hyperinsulinemia. Cancer Discov 14, 240–257. 10.1158/2159-8290.CD-23-0944

von Willebrand, M., Williams, S., Saxena, M., Gilman, J., Tailor, P., Jascur, T., Amarante-Mendes, G.P., Green, D.R., Mustelin, T., 1998. Modification of phosphatidylinositol 3-kinase SH2 domain binding properties by Abl-or Lck-mediated tyrosine phosphorylation at Tyr-688. J. Biol. Chem. 273, 3994–4000. 10.1074/jbc.273.7.3994

Walser, R., Burke, J.E., Gogvadze, E., Bohnacker, T., Zhang, X., Hess, D., Küenzi, P., Leitges, M., Hirsch, E., Williams, R.L., Laffargue, M., Wymann, M.P., 2013. PKCβ phosphorylates PI3Kγ to activate it and release it from GPCR control. PLoS Biol. 11, e1001587. 10.1371/journal.pbio.1001587

Yu, J., Zhang, Y., McIlroy, J., Rordorf-Nikolic, T., Orr, G.A., Backer, J.M., 1998. Regulation of the p85/p110 phosphatidylinositol 3’-kinase: stabilization and inhibition of the p110alpha catalytic subunit by the p85 regulatory subunit. Mol Cell Biol 18, 1379– 1387. 10.1128/MCB.18.3.1379

